# Effect of a Sepsis Prediction Algorithm on Patient Mortality, Length of Stay, and Readmission

**DOI:** 10.1101/457465

**Authors:** Hoyt Burdick, Eduardo Pino, Denise Gabel-Comeau, Andrea McCoy, Carol Gu, Jonathan Roberts, Joseph Slote, Nicholas Saber, Jana Hoffman, Ritankar Das

**Affiliations:** Cabell Huntington Hospital, Huntington, WV; Marshall University School of Medicine, Huntington, WV; Cape Regional Medical Center, Cape May Court House, New Jersey, USA; Dascena, Inc., Hayward, CA, United States

## Abstract

**Objective:** To validate performance of a machine learning algorithm for severe sepsis determination up to 48 hours before onset, and to evaluate the effect of the algorithm on in-hospital mortality, hospital length of stay, and 30-day readmission.

**Setting:** This cohort study includes a combined retrospective analysis and clinical outcomes evaluation: a dataset containing 510,497 patient encounters from 461 United States health centers for retrospective analysis, and a multiyear, multicenter clinical data set of real-world data containing 75,147 patient encounters from nine hospitals for clinical outcomes evaluation.

**Participants:** For retrospective analysis, 270,438 adult patients with at least one documented measurement of five out of six vital sign measurements were included. For clinical outcomes analysis, 17,758 adult patients who met two or more Systemic Inflammatory Response Syndrome (SIRS) criteria at any point during their stay were included.

**Results:** At severe sepsis onset, the MLA demonstrated an AUROC of 0.91 (95% CI 0.90, 0.92), which exceeded those of MEWS (0.71, P<001), SOFA (0.74; P<.001), and SIRS (0.62; P<.001). For severe sepsis prediction 48 hours in advance of onset, the MLA achieved an AUROC of 0.77 (95% CI 0.73, 0.80). For the clinical outcomes study, when using the MLA, hospitals saw an average 39.5% reduction of in-hospital mortality, a 32.3% reduction in hospital length of stay, and a 22.7% reduction in 30-day readmission rate.

**Conclusions:** The MLA accurately predicts severe sepsis onset up to 48 hours in advance using only readily available vital signs in retrospective validation. Reductions of in-hospital mortality, hospital length of stay, and 30-day readmissions were observed in real-world clinical use of the MLA. Results suggest this system may improve severe sepsis detection and patient outcomes over the use of rules-based sepsis detection systems.

**KEY POINTS:** *Question:* Is a machine learning algorithm capable of accurate severe sepsis prediction, and does its clinical implementation improve patient mortality rates, hospital length of stay, and 30-day readmission rates?

*Findings:* In a retrospective analysis that included datasets containing a total of 585,644 patient encounters from 461 hospitals, the machine learning algorithm demonstrated an AUROC of 0.93 at time of severe sepsis onset, which exceeded those of MEWS (0.71), SOFA (0.74), and SIRS (0.62); and an AUROC of 0.77 for severe sepsis prediction 48 hours in advance of onset. In an analysis of real-world data from nine hospitals across 75,147 patient encounters, use of the machine learning algorithm was associated with a 39.5% reduction in in-hospital mortality, a 32.3% reduction in hospital length of stay, and a 22.7% reduction in 30-day readmission rate.

*Meaning:* The accurate and predictive nature of this algorithm may encourage early recognition of patients trending toward severe sepsis, and therefore improve sepsis related outcomes.

**STRENGTHS AND LIMITATIONS OF THIS STUDY:**

- A retrospective study of machine learning severe sepsis prediction from a dataset with 510,497 patient encounters demonstrates high accuracy up to 48 hours prior to onset.
- A multicenter clinical study of real-world data using this machine learning algorithm for severe sepsis alerts achieved reductions of in-hospital mortality, length of stay, and 30-day readmissions.
- The required presence of an ICD-9 code to classify a patient as severely septic in our retrospective analysis potentially limits our ability to accurately classify all patients.
- Only adults in US hospitals were included in this study.
- For the real-world section of the study, we cannot eliminate the possibility that implementation of a sepsis algorithm raised general awareness of sepsis within a hospital, which may lead to higher recognition of septic patients, independent of algorithm performance.

## Introduction

Severe sepsis and septic shock are a dysregulated response to infection, and they are among the leading causes of death in the United States. Approximately 750,000 patients are diagnosed with severe sepsis annually, with a high associated mortality [1–2]. The cost of treating sepsis is estimated to be $16.7 billion per year, making sepsis one of the most expensive conditions to diagnose and treat [2,6–3].

Despite this, sepsis is difficult to detect and predict. New definitions intended to improve the clinical recognition of sepsis have recently been proposed [3–4,5]. The previous use of screening based on Systemic Inflammatory Response Syndrome (SIRS) criteria was found to be nonspecific [76]. Multiple studies have shown that accurate early diagnosis and treatment, including sepsis bundle compliance, can reduce the risk of adverse patient outcome from severe sepsis and septic shock [8,7–9]. Therefore, earlier detection and more accurate recognition of patients at high risk of developing severe sepsis or septic shock provide a valuable window for effective sepsis treatments.

In this study, we evaluated the performance of our machine learning algorithm (MLA) for sepsis prediction and detection. The MLA determines risk of sepsis using data from patient Electronic Health Records. We evaluated the performance metrics of the algorithm including Area Under the Receiver Operating Curve (AUROC), sensitivity, and specificity, using both retrospective patient data from 461 hospitals and real world data from nine diverse clinical settings. In our retrospective study, we compared AUROC values of the MLA to AUROC values obtained by other standard scoring systems. In our clinical outcomes analysis, we collected real-world patient data from nine hospitals using the MLA and evaluated the effect of the algorithm on in-hospital patient mortality, hospital length of stay, and 30-day readmissions. The MLA evolved over time, as more data became available to train it. Thus, the real-world data was not based a single static algorithm, but rather a single algorithmic design. The latest state of the MLA was characterized in the retrospective analysis. Previous states of the algorithm have been studied retrospectively and prospectively [10–16]; however, this study was performed on significantly larger and more diverse datasets.

## Methods

### Dataset

The Dascena Analysis Dataset (DAD) and the Cabell Huntington Hospital Dataset (CHHD) were used for all experiments for retrospective algorithm validation. The DAD is comprised of 489,850 randomly-selected inpatient and emergency department encounters obtained from de-identified EHR records at 461 total academic and community hospitals across the continental United States (Supplementary Table 1). Data were collected between 2001 and 2015, with the majority of encounters occurring between 2014 and 2015. Details about all hospitals are provided in Supplementary Table 2. The CHHD includes 20,647 inpatient and emergency encounters from Cabell Huntington Hospital (Huntington, WV) collected during 2017.

Prospectively collected real-world patient data was abstracted from the Electronic Health Record (EHR) systems of Epic (Epic Systems, Verona, Wisconsin, USA), Allscripts (Allscripts Healthcare Solutions, Chicago, Illinois, USA), Cerner (Cerner Systems, North Kansas City, Missouri, USA), Meditech (Meditech, Westwood, Massachusetts, USA), Paragon (McKesson Corporation, San Francisco, California, USA), and Soarian (Cerner Systems, North Kansas City, Missouri, USA), across the nine hospitals in the clinical outcomes evaluation.This data spanned 75,147 patient encounters from between early 2017 to mid-2018. Details about these nine hospitals are also provided in Supplementary Table 1.

All patient information was de-identified prior to analysis in compliance with the Health Insurance Portability and Accountability Act (HIPAA). Data collection for all datasets was passive and did not impact patient safety. Approval for this study was exempted by the Institutional Review Board (IRB) at Pearl Pathways.

### Patient measurements and inclusion criteria

In both our retrospective and clinical outcomes analyses, we analyzed only adult EHR records (ages 18 and over) from inpatient (including critical care) wards and emergency departments. All genders and ethnicities were included.

For inclusion in the retrospective analysis, patient records must have contained at least one documented measurement of five out of six vital sign measurements including heart rate, respiration rate, temperature, diastolic and systolic blood pressure, and SpO_2_. We also required at least one recorded observation of each measurement required to calculate the SOFA score, including Glasgow Coma Scale, PaO_2_/FiO_2_, bilirubin level, platelet counts, creatinine level, and mean arterial blood pressure or administration of vasopressors (see Calculating Comparators below). All patients who presented with sepsis on admission were excluded. These criteria resulted in the inclusion of 270,438 patients from the DAD (See Supplementary Table 3). Patients were divided into subgroups based on hospital length of stay in order to assess MLA performance at each predetermined prediction time. Patients were included in analysis only if their length of stay exceeded the tested prediction time. This resulted in decreasing subgroup size as prediction time was increased. For each prediction time, patients who became severely septic within two hours of the prediction window were excluded. This ensured the presence of adequate data with which to train and test the algorithm for each prediction task. For any patient with a stay exceeding 2,000 hours, the last 2,000 hours of hospital data were used for the study, in order to limit the size of data analysis matrices.

For the clinical outcomes analysis, patient stays that met two or more Systemic Inflammatory Response Syndrome (SIRS) criteria at any point during their stay were considered "sepsis-related” and included. This resulted in the inclusion of 17,758 patient encounters for analysis.

### Data Fields

In this study, demographic, admission and discharge times, vital sign, laboratory, and drug administration data were abstracted, for each visit of a given patient, from the applicable dataset for each of retrospective and clinical outcomes analyses. Not all fields were available at all facilities. Details on which data fields were abstracted are included in Supplementary Table 4.

### Binning and Imputation

For retrospective analysis, vital sign and laboratory measurements were binned by the hour for each included patient, beginning at the time of the patient’s first recorded measurement and ending with the last whole hour of available data observed before the patient’s final measurement. Predictions for the MLA were made using only systolic blood pressure, diastolic blood pressure, heart rate, temperature, respiratory rate and SpO_2_ measurements. If one of the required measurements was not recorded for a given hour, the missing measurement was filled with the most recently recorded previous measurement of that type (i.e. carry-forward imputation). If multiple measurements of a single type were recorded within a given hour, these measurements were averaged to produce a single value for the hour, thus minimizing information fed to the classifier regarding sampling frequency. All data processing was done using Python software [17].

### Gold Standard

For retrospective analysis, we defined our severe sepsis gold standard as “organ dysfunction caused by sepsis,” with sepsis defined as “the presence of two or more SIRS criteria paired with a suspicion of infection” [3]. To positively identify cases of severe sepsis, we conservatively required the presence of an International Classification of Diseases (ICD) 9 code 995.9x. We defined onset time as the first time at which two SIRS criteria and at least one organ dysfunction criteria (Supplementary Table 5) were met within the same hour.

### Calculating Comparators

In the retrospective analysis, we fixed sepsis identification score thresholds of 2, 2, and 1 for MEWS, SOFA, and SIRS criteria, respectively. In other words, a MEWS score ≥ 2 indicates a patient would be categorized by MEWS as septic; all such comparators were calculated for sepsis detection at the time of onset assigned by the gold standard. Because the MLA can obtain a range of sensitivities and specificities, to report the MLA sensitivity and specificity we selected a fixed point on the Receiver Operating Characteristic (ROC) curve with sensitivity near 0.80. This facilitated table-based comparisons of specificity while holding sensitivity relatively constant.

We compared performance of the MLA and rules-based systems using the area under the receiver operating characteristic (AUROC) curve.

### The Machine Learning Algorithm

We constructed our classifier using gradient boosted trees, implemented in Python (Python Software Foundation, https://www.python.org/), with the XGBoost package [18]. Predictions were generated from the binned values for the vital signs of systolic blood pressure, diastolic blood pressure, heart rate, temperature, respiratory rate and SpO_2_ at prediction time, one hour before prediction time, and two hours before prediction time, as well as the hourly differences between each of these measurements. These values were concatenated into a feature vector with fifteen elements. The gradient boosted trees approach builds an ensemble of decision trees. The ensemble then makes a prediction based on an aggregate of these scores.

Tree branching was determined by discretizing vital sign measurements into two categories, and patient risk scores were determined by their final categorization in each tree. We limited tree branching to six levels, included no more than 1000 trees in the final ensemble, and set the XGBoost learning rate to 0.1. These hyperparameters were chosen to align with previous work and justified in the context of the present data with a coarse grid search [10].

### Study Design

For retrospective analysis, model performance was validated using ten-fold cross validation. To perform ten-fold cross validation, we first randomly shuffled the DAD dataset and divided it into two portions: 80% of the dataset was used as a training set, and 20% of the dataset was withheld as an independent testing set. We then further divided the training set into tenths, training the algorithm on nine of these tenths and assessing its performance on the remaining tenth. We repeated this process ten times, using each possible combination of training and testing folds. We then assessed each of the resulting ten models on the independent testing set. Reported performance metrics on the testing set are the average performance of each of these ten models on the testing set. The reported threshold score, the MLA score at which a patient was deemed to be positive for severe sepsis, was taken from the best performing model from each ten-fold cross validation, with performance measured by AUROC value.

For clinical outcomes analysis, we collected data from nine hospitals that implemented the MLA for sepsis prediction and detection. We then evaluated that data to determine the effect the algorithm had on patient outcomes of in-hospital mortality, hospital length of stay, and 30-day readmission. Providers at the hospitals using the MLA received telephonic alerts if the MLA score was above a threshold set by the hospital.

Patients were considered to be “sepsis-related” and included for analysis if they met two or more SIRS criteria at any point during their stay in units where the MLA was used and were over the age of 18. We classified patients in this manner due to the predictive nature of the MLA. Because the algorithm is designed to identify patients likely to develop sepsis, including only those patients who met the 2001 consensus severe sepsis or septic shock definition criteria or the more recent Sepsis-3 criteria may have excluded patients who would have developed sepsis had they not been identified and treated early. The SIRS criteria, while non-specific, are associated with early sepsis diagnostic criteria, and their use in this study ensured that those patients most at risk for sepsis were included in our final analysis.

Telephonic notification volumes differed from month to month during the trial period. Months with uncharacteristically low volumes (fewer than 5) were excluded from analysis. Three sites in the study were affected by the exclusion of certain months from analysis. Alert volumes varied as site-specific customization was performed through PDSA (plan-do-study-act) cycles for thresholding and rules-based suppression to optimize the algorithm for the best fit into a given care setting. When data from the period preceding implementation of the MLA was not available the baseline period used was the month immediately following implementation. This was the case for five out of the nine hospitals. The analysis was repeated using three of the nine hospitals which had at least one month of baseline data preceding MLA implementation, and the outcomes were similar. Not all three patient metrics were measured at all sites. Length of stay was measured at all sites, in-hospital mortality was measured at six out of nine of the sites, and readmission was measured at five out of nine sites. If admission and discharge time stamps were unavailable, length of stay and readmission were determined by defining new visits when vital sign measurements for a given patient were observed to be greater than 120 hours apart.

### Statistical Tests

Two-sample *t*-tests were used to determine if there was a statistically significant difference in means between the baseline and MLA periods for sepsis-related LOS. We used the two-proportion risk difference z-test to determine if there was a statistically significant decrease of the in-hospital mortality or the 30-day readmission rate with the use of the MLA. All tests were two-tailed with an alpha level of 0.05, and were performed using Python.

## Results

The MLA demonstrated a higher severe sepsis detection AUROC (0.91; 95% confidence interval (CI) 0.900,0. 920) than MEWS (0.71; P<.001), SOFA (0.74; P<.001), and SIRS (0.62; P<.001) (Figure 1). Detailed performance metrics for all scoring systems at the time of severe sepsis onset are presented in Table 2.

**Table 1:** Demographics Table. Demographic and clinical characteristics of patients included in the Dascena Analysis Dataset and for clinical outcomes analysis.

|  | Retrospective Analysis |  | Clinical Outcomes Analysis |  |
| --- | --- | --- | --- | --- |
|  | Septic | Non-Septic | Baseline | MLA |
| Total Number | 20,876 | 468,974 | 12,793 | 62,354 |
| Age (SD) | 62 (17.0) | 56 (19.0) | 45 (24.4) | 45 (24.0) |
| Male | 10,326 (49.5%) | 221,029 (45.1%) | 5,429 (42.4%) | 26,126 (41.9%) |
| Female | 9,325 (44.7%) | 219,866 (44.9%) | 7,364 (57.6%) | 36,228 (58.1%) |
| Unknown | 1,225 (5.9%) | 28,079 (6.0%) | — | — |
| White | 10,257 (49.1%) | 165,875 (33.9%) | 11,832 (87.7%) | 54,635 (81.8%) |
| Black | 1,267 (6.1%) | 23,734 (5.1%) | 594 (4.4%) | 2,469 (3.7%) |
| Hispanic | 1,090 (5.22%) | 33,944 (7.24%) | 1,063 (7.87%) | 9,609 (14.4%) |
| Asian American | 355 (1.7%) | 4,953 (1.1%) | 10 (0.1%) | 44 (0.1%) |
| Unknown | 7,907 (37.9%) | 240,468 (49.1%) | — | — |
| Temperature | 36.9 (0.3) | 36.8 (0.2) | 36.8 (0.3) | 36.8 (0.3) |
| Respiratory Rate | 20.8 (7.5) | 18.1 (5.1) | 18.2 (4.7) | 18.2 (4.1) |
| Systolic Blood Pressure | 119.1 (16.7) | 125.5 (16.7) | 127.0 (18.0) | 129.5 (19.1) |
| Diastolic Blood Pressure | 66.6 (9.8) | 73.1 (10.5) | 72.9 (11.0) | 75.1 (11.5) |
| Heart Rate | 93.8 (16.7) | 85.4 (17.1) | 84.7 (17.3) | 86.1 (18.5) |
| Lactate | 2.63 (1.9) | 1.90 (1.6) | 1.9 (1.70) | 1.9 (1.86) |
| Creatinine | 1.7 (1.8) | 1.5 (2.6) | 1.4 (2.54) | 1.2 (1.70) |
| International Normalized Ratio (INR) | 1.4 (0.8) | 1.1 (0.4) | 1.2 (0.58) | 1.3 (0.90) |
| Platelets | 238.5 (105.1) | 239.6 (75.1) | 239.5 (77.1) | 241.9 (85.0) |
| SpO<sub>2</sub> | 96.8 (1.5) | 97.6 (1.3) | 97.4 (1.6) | 97.4 (1.7) |
| White Blood Count | 8.5 (1.44) | 8.2 (1.72) | 8.4 (2.51) | 8.2 (2.03) |
| PaO<sub>2</sub> | 95.6 (27.8) | 102.6 (45.3) | 101.7 (47.9) | 103.8 (51.8) |
| Bilirubin | 1.3 (2.5) | 0.7 (1.2) | 0.7 (1.3) | 0.7 (1.1) |
| FiO<sub>2</sub> | 47.4 (20.4) | 42.0 (18.1) | 44.3 (20.4) | 46.6 (22.2) |
| pH | 7.4 (0.09) | 7.4 (0.08) | 7.4 (0.08) | 7.4 (0.09) |

**Table 2:** Comparison Table of Performance Metrics for MLA to Standard Scoring Systems, at Onset. Detailed performance metrics for the Machine Learning Algorithm (MLA) and rules-based systems taken at time of severe sepsis onset. The score threshold reported for the MLA is the average over rounds of 10-fold cross-validation. AUROC: Area Under the Receiver Operating Characteristic; MEWS: Modified Early Warning Score; SOFA: Sequential Organ Failure Assessment; SIRS: Systemic Inflammatory Response Syndrome; DOR: Diagnostic Odds Ratio; LR: Likelihood Ratio; F1 Score: the harmonic average of the precision and recall; PPV: Positive Predictive Value; NPV: Negative Predictive Value.

|  | MLA ≥ 4.05 | MEWS ≥ 2 | SOFA ≥ 2 | SIRS ≥ 1 |
| --- | --- | --- | --- | --- |
| **AUROC (95% CI)** | 0.91<br>(0.900, 0.920) | 0.705<br>(0.696, 0.713) | 0.744<br>(0.737, 0.750) | 0.623<br>(0.615, 0.632) |
| **Sensitivity** | 0.801 | 0.834 | 0.787 | 0.839 |
| **Specificity** | 0.867 | 0.448 | 0.554 | 0.336 |
| **Accuracy** | 0.937 | 0.609 | 0.647 | 0.65 |
| **DOR** | 25.812 | 4.111 | 4.636 | 2.684 |
| **LR+** | 7.946 | 1.51 | 1.764 | 1.264 |
| **LR-** | 0.231 | 0.372 | 0.384 | 0.478 |
| **F1 Score** | 0.295 | 0.046 | 0.052 | 0.042 |
| **PPV** | 0.192 | 0.024 | 0.027 | 0.022 |
| **NPV** | 0.999 | 0.999 | 0.999 | 0.999 |

**Figure 1:**
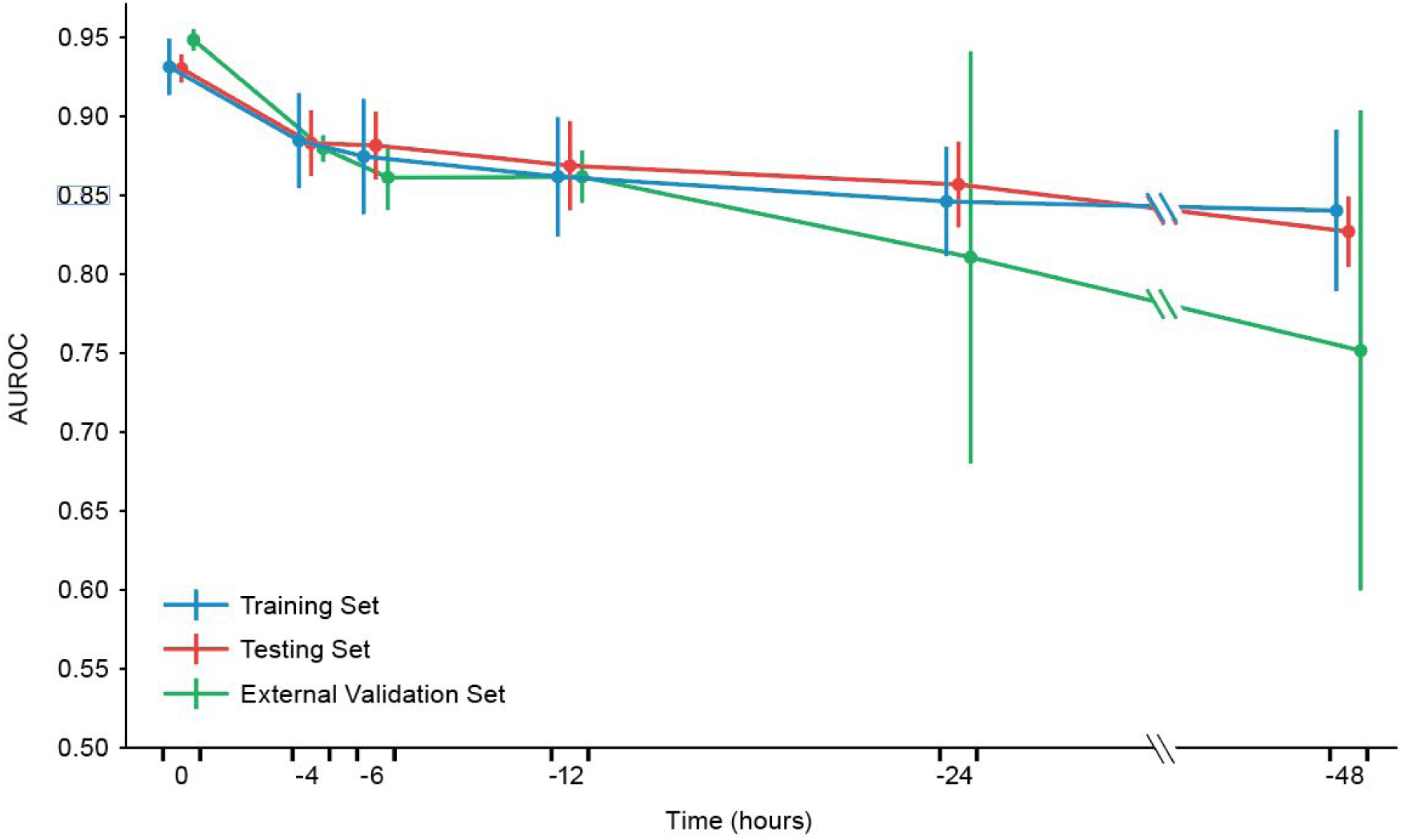
AUROC Over Time. Depicts performance of the MLA in predicting the onset of severe sepsis at 0, 2, 4, 6, 12, 24 and 48 hours before severe sepsis onset. "Training” and "Testing” Set were derived from application on the Dascena Analysis Dataset, and the "External Validation Set” was derived from application on the Cabell Huntington Hospital Dataset.

**Figure 2:**
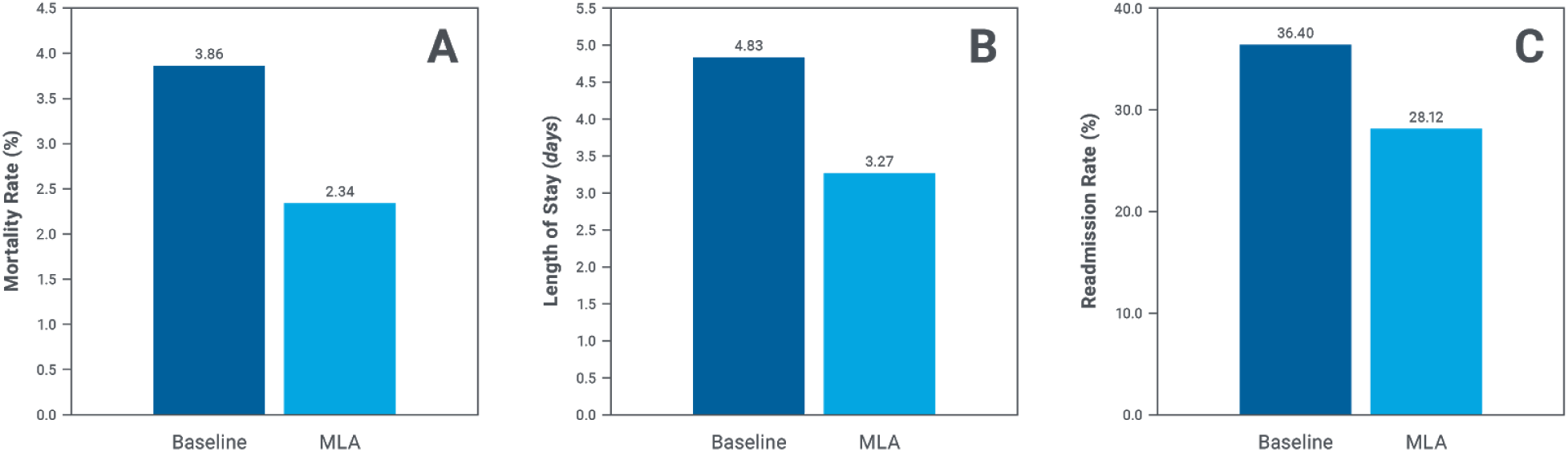
Patient Outcomes on External Validation Dataset. Differences in (A) in-hospital mortality, (B) hospital length of stay (LOS), and (C) 30-day readmissions in the baseline period and the MLA period. Use of the MLA was associated with a 39.5% reduction of in-hospital mortality (P<.001), a 32.3% reduction in LOS (P<.001), and a 22.7% reduction in thirty-day readmissions (P<.001).

Additionally, the MLA maintained a superior AUROC for all prediction windows as compared to all onset-time rules-based scoring systems; at 48 hours prior to severe sepsis onset, the MLA demonstrated an AUROC value of 0.77 (Figure 1). Detailed performance metrics for the MLA at all prediction windows are presented in Supplementary Table 6.

We ranked feature importance for severe sepsis detection and prediction using the MLA using average entropy gain for each feature. Feature importance varied significantly by prediction window (Supplementary Figure 1).

For clinical outcomes analysis, we collected data from nine hospitals that implemented the MLA for sepsis prediction and detection. Providers at the hospitals using the MLA received telephonic alerts if the MLA score was above a threshold set by the hospital. The outcomes after MLA implementation were a 39.50% reduction of in-hospital mortality, a 32.27% reduction of length of stay, and a 22.74% reduction in 30-day readmission (Table 4). The analysis was repeated for a subset of three hospitals with at least one month of baseline (pre-MLA implementation) data, with a total of 52,487 patients. This resulted in 3,951 patients in the baseline period (971 included as sepsis-related, see above), and 48,536 patients (10,646 included as sepsis-related) in the MLA analysis period. The outcomes for this patient subset were a 42.50% reduction of in-hospital mortality and a 23.82% reduction in length of stay.

**Table 4:** Patient Outcomes Table. Analysis of in-hospital mortality, hospital length of stay, and 30-day readmissions, in the baseline and MLA periods. There were 12,793 patients in the baseline period, of whom, 3,592 were included for analysis and 62,354 patients in the MLA period, of whom, 14,166 patients were included for analysis.

|  | Baseline Period | MLA Period | Reduction |
| --- | --- | --- | --- |
| In-Hospital Mortality | 3.86% | 2.34% | 39.50% |
| Length of Stay | 4.83 days | 3.27 days | 32.27% |
| 30-Day Readmission | 36.4% | 28.12% | 22.74% |

## Discussion

The machine learning algorithm more accurately detected hospital acquired severe sepsis onset than the commonly used rules-based disease severity scoring systems MEWS, SOFA, and SIRS. Up to 48 hours before onset, the MLA maintained high AUROC values and demonstrated higher sensitivity and specificity than commonly used rules-based systems applied at the time of onset. The accuracy of the MLA, paired with the minimal patient data required for predictions, suggests that this system may improve severe sepsis detection and patient outcomes over the use of a rules-based system. The high specificity of the MLA may help to reduce alarm fatigue, a known patient safety hazard [20].

In this study, the algorithm was tested on a large and diverse retrospective dataset containing inpatient and emergency department patient data from 461 teaching and non-teaching hospitals in the US. This dataset includes patient data from intensive care unit and floor wards, representing a variety of data collection frequencies and care provision levels. Sepsis manifestation can vary depending on factors such as patient race and comorbidities [21]; it is therefore important that sepsis detection methods be validated across a diverse population in order to ensure accurate discrimination for all patients. The high performance of the MLA on this dataset indicates that this algorithm may be able to improve patient outcomes in a variety of clinical settings. In addition to strong performance on a hold-out test set, consistent performance on an external validation set reduces chances that the reported results are caused by over fit.

In the clinical outcomes analysis, the MLA was associated with reductions of in-hospital mortality, hospital length-of-stay and 30-day hospital readmissions. The demonstrated reduction in average length of stay provides a potential financial impact for the hospitals where MLA was implemented. The average length of stay reduction was found to be 1.56 days. At an average per diem cost of care of $2,271, the reduction of length of stay translates to approximately $14.5 million of annual cost savings across all nine hospitals included in this analysis. These findings on post-marketing real-world data confirm pre-marketing randomized clinical trial results [11]. The MLA system used in both analyses demonstrated higher sensitivity and specificity than comparable rule-based systems (MEWS, SOFA, and SIRS), and accuracy of the MLA suggests that this system may improve severe sepsis detection and patient outcomes over the use of comparators. It is worth noting that the standard deviation for the external validation dataset — which quantifies variability in patient populations [22] — becomes larger at longer look-ahead times. This indicates increased variation in the performance of the algorithm at longer look-ahead times in the external validation set.

Previous research has shown that early detection and treatment of sepsis can improve patient outcomes [8,9,23]. The long prediction times provided by this algorithm may encourage early recognition of patients trending toward severe sepsis, and therefore improve sepsis related outcomes.

## Limitations

While the retrospective analysis incorporated data from a large number of institutions (nearly 10% of US hospitals), we cannot claim generalizability to additional specific settings or populations on the basis of this study. Generalizability of the retrospective results is also limited by our inclusion criteria requiring that all patients manifesting severe sepsis within two hours of each prediction window be excluded from the analysis.

The required presence of an ICD-9 code to classify a patient as severely septic in our retrospective analysis potentially limits our ability to accurately classify all patients in the dataset [24]. However, past research has shown ICD-9 coding to be a reasonable means of retrospectively detecting patients with severe sepsis [25,26]. Further, our gold standard criteria may also limit the accuracy of our severe sepsis onset time analysis, as the time a condition was recorded in the patient chart may not represent the time the condition actually manifested.

In the retrospective analysis, we treated severe sepsis detection and prediction as a classification task. While a time-to-event modeling approach would have also been possible, classification methods are significantly more common in the literature [19, 27–30]. By using the same modelling approach, the present study can be readily compared with existing work on sepsis detection models using standard metrics such as AUROC and specificity.

Our real-world data analysis has several limitations. First, clinician and team responses to patients at possible risk for sepsis vary by hospital. Second, only adults in US hospitals were included in the study. While nine diverse hospitals were included in the analysis, these hospitals may not be representative of all US hospitals. Third, baseline data were not available for all hospitals and the first month of MLA data was used as an approximation in these cases. This may lead to an underestimation of the effect of the MLA at these sites. Additionally, data was not available from all hospitals for all months and outcome measurements. However, the analysis was repeated on a subset of three hospitals with at least one month of baseline pre-MLA implementation data and outcomes were similar. Fourth, the study did not follow patient mortality after hospital discharge. Fifth, we cannot eliminate the possibility that implementation of a sepsis algorithm raised general awareness of sepsis within a hospital, which may lead to higher recognition of septic patients, independent of algorithm performance.

## Conclusion

This study validates a machine learning algorithm for severe sepsis detection and prediction against a diverse retrospective dataset containing patient data from 461 US hospitals. The algorithm is capable of predicting severe sepsis onset up to 48 hours in advance of onset using only six frequently collected patient measurements, and demonstrates higher sensitivity and specificity than commonly used sepsis detection methods such as MEWS, SOFA and SIRS.

In a clinical outcomes analysis of the algorithm across nine hospitals, use of the MLA was associated with a 39.5% reduction of in-hospital mortality, a 32.3% reduction in hospital length of stay, and a 22.7% reduction in 30-day readmissions. These results support that the implementation of an accurate machine learning algorithm for early sepsis recognition may lead to improved patient outcomes.

## Acknowledgments

We gratefully acknowledge Yvonne Zhou for assistance with data analysis, and Touran Fardeen and Emily Pellegrini for assistance with manuscript editing.

## Contributors

RD conceived the described experiments. HB and EP acquired the Cabell Huntington Hospital (CHH) data. JR, JS, and NS executed the experiments. RD, JR, JS, NS and JH interpreted the results. RD and JH wrote the manuscript. HB, EP, DGC, AM, CG, JR, JS, NS, JH, and RD revised the manuscript.

## Disclaimer

The content is solely the responsibility of the authors and does not necessarily represent the official views of the National Science Foundation.

## Patient Consent

Not required.

## Provenance and Peer Review

Not commissioned; externally peer reviewed.

## Data Sharing Statement

No data obtained from Cabell Huntington Hospital in this study can be shared or made available for open access.

## Open Access

This is an Open Access article distributed in accordance with the Creative Commons Attribution Non Commercial (CC BY-NC 4.0) license, which permits others to distribute, remix, adapt, build upon this work non-commercially, and license their derivative works on different terms, provided the original work is properly cited and the use is non-commercial. See: http://creativecommons.org/licenses/by-nc/4.0/

